# Benefits of spaced learning are predicted by re-encoding of past experience in ventromedial prefrontal cortex

**DOI:** 10.1101/2024.05.14.594263

**Authors:** Futing Zou, Brice A. Kuhl, Sarah DuBrow, J. Benjamin Hutchinson

## Abstract

More than a century of research shows that spaced learning improves long-term memory. Yet, there remains debate concerning why. A major limitation to resolving theoretical debates is the lack of evidence for how neural representations change as a function of spacing. Here, leveraging a massive-scale 7T human fMRI dataset, we tracked neural representations and behavioral expressions of memory as participants viewed thousands of natural scene images that repeated at lags ranging from seconds to many months. We show that spaced learning increases the similarity of human ventromedial prefrontal cortex representations across stimulus encounters and, critically, these increases parallel and predict the behavioral benefits of spacing. Additionally, we show that these spacing benefits critically depend on remembering and, in turn, ‘re-encoding’ past experience. Collectively, our findings provide fundamental insight into how spaced learning influences neural representations and why spacing is beneficial.

---

One of the most robust and well-documented phenomena in human memory research is that when repetitions of a stimulus are spaced over time—as opposed to massed—long-term memory for that stimulus is improved^1^. This spacing effect operates across a wide variety of memoranda^2^ and species^3^ and also across impressively long timescales^4–7^. However, despite the ubiquity of this effect and its significant implications for memory in everyday and educational settings, there remains debate concerning why spaced learning improves memory. Why is a repeated stimulus more effectively encoded when more time has elapsed since it was previously encoded?

While a number of theoretical accounts of spacing effects have been advanced^3,6,8^, one of the most prominent ideas is that spaced learning benefits memory by increasing encoding variability^9–11^. By this account, when a stimulus is re-encountered after a long delay, it is encoded *differently* than the first time it was encountered. A more variable representation is argued to have more ‘points of access’ and, therefore, to increase the likelihood of later retrieval^12^. However, if encoding variability were the only factor, memory would monotonically increase with greater spacing (the more spacing, the better). In contrast, studies have shown that as the lag between stimulus repetitions increases, the benefit of spacing will eventually reverse, producing an ‘inverted U’ shaped relationship between spacing and subsequent memory^2,4,6^. This non-monotonic pattern is often only observed when long timescales are considered (spacing on the order of weeks or months), but it is extremely informative, at a theoretical level, because it suggests a second factor also contributes to spacing effects—a factor that explains diminished benefits at very long lags. Perhaps the most commonly-advanced second factor is study-phase retrieval^13,14^. By this account, when a stimulus is re-encountered, it triggers retrieval of the original encounter (benefitting memory), but the benefit of retrieval is negatively related to lag because forgetting of the first encounter becomes more likely with longer lags. While the combination of encoding variability and study-phase retrieval has high explanatory power^6,10,12,15–17^, a fundamental limitation of almost all leading theoretical accounts of spacing effects is that they are not directly informed, or constrained, by experimental evidence of how spacing influences neural representations. This is particularly glaring in the case of encoding variability, which makes obvious predictions about the variability of neural representations over time and the relationship of this variability to subsequent memory.

The lack of integration of neural evidence with cognitive theories of spacing effects is partly due to the surprisingly limited number of neuroimaging / electrophysiological studies that have characterized neural representations as a function of spacing. Moreover, existing evidence is largely limited to studies that measure spacing effects over short timescales (a single experimental session), making it difficult to test ideas from cognitive theories that are specifically motivated by behavioral spacing effects over long timescales. A primary issue is that it is difficult, or at least resource intensive, to measure neural representations across days, weeks, or months. That said, there is recent evidence that, at least at short timescales, neural representations of repeated events may actually be *more similar* with greater spacing^18,19^—a finding not predicted or easily explained by an encoding-variability account. However, it is currently unclear why neural representations would become more similar with spacing, how neural similarity might change over longer timescales, and what neural similarity ultimately tells us about why spaced learning is beneficial.

Here, we measured neural representations and behavioral expressions of memory for stimuli repeated over long timescales with the goal of integrating neural and behavioral measures to inform theories of spacing effects. To this end, we leveraged the Natural Scenes Dataset^20^—an unprecedented dataset that combines behavioral measures of memory and 7T fMRI for human participants that studied stimuli distributed across 30-40 experimental sessions over an 8-10 month window (**Fig. 1a-b**). Across these sessions, participants viewed 9,209-10,000 natural scene images presented up to three times each (E1, E2, E3) and made recognition memory decisions at each encounter (continuous recognition task). This yielded a remarkably wide range of lags (spacing) between the first two exposures, from 4 seconds up to 302 days (**Fig. 1c**). Of central interest here was how spacing between the first two exposures with a stimulus (E1-E2 lag) influenced corresponding neural similarity (E1-E2 similarity) and whether neural measures of similarity parallel and predict behavioral benefits of spacing (recognition memory at E3). We focused our analyses on ventromedial prefrontal cortex (vmPFC), motivated by the extensive human neuroimaging literature implicating vmPFC in episodic memory across long timescales^21–24^ and by recent evidence of spacing effects on neural similarity in the rodent medial prefrontal cortex^19^. To preview, we show spacing-related increases in vmPFC similarity that (a) parallel and predict behavioral benefits of spacing, and (b) reflect the re-encoding of memories for prior encounters.

**Fig. 1:**
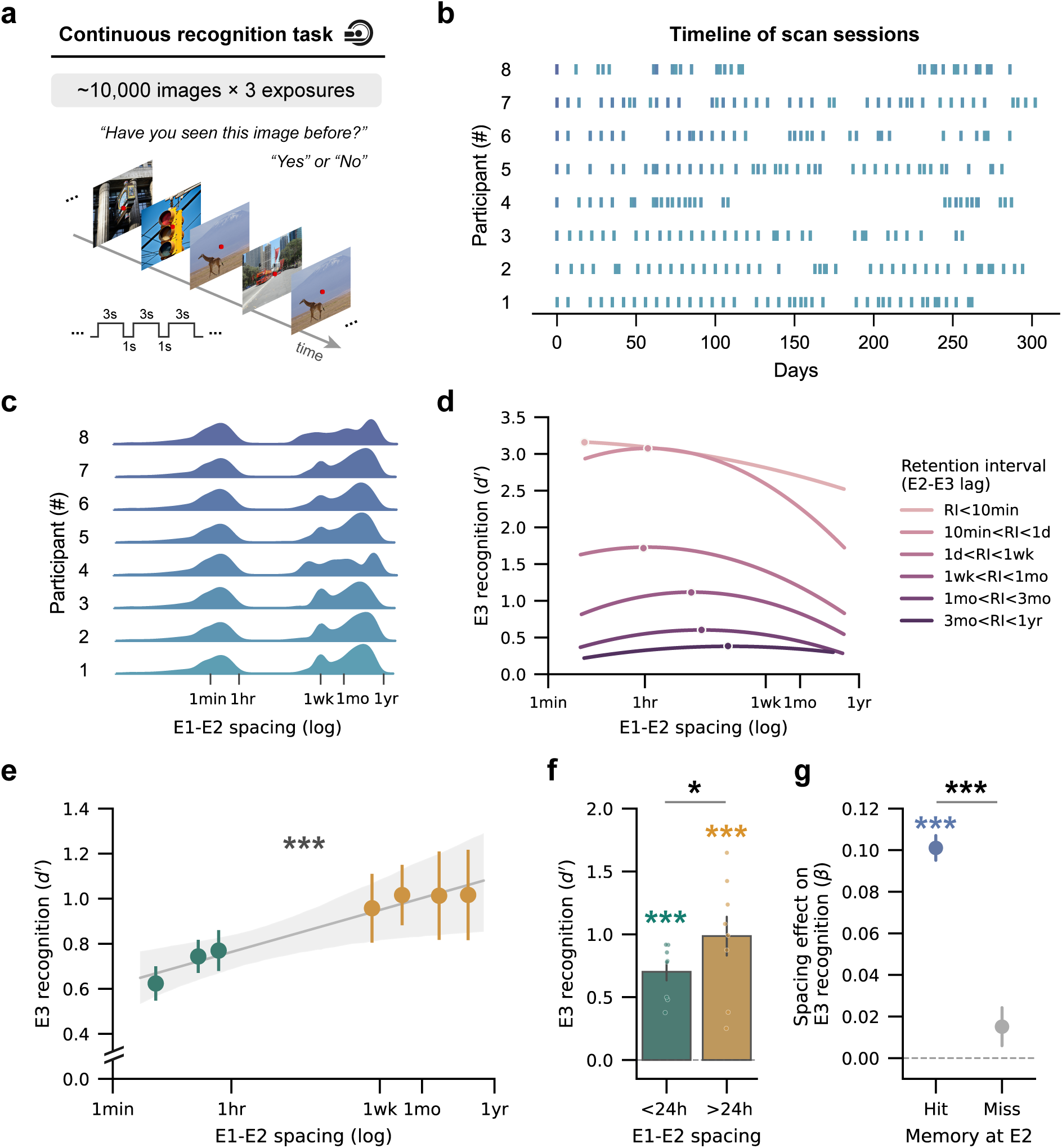
Task and behavioral results. **a,** Trial structure. During fMRI, participants performed a continuous recognition task in which thousands of natural scene images were presented up to three times each with a pseudo-random delay interval between repetitions. **b,** Timeline of 7T fMRI scan sessions. Each of the eight participants completed 30-40 scan sessions over a 10-month window. The first scan session for each participant corresponds to day 0. **c,** Smoothed histograms showing, for each participant, the relative frequencies of various E1-E2 spacing intervals (from seconds to months), shown on a log scale. **d,** Relationships between spacing (E1-E2 lag) and subsequent memory (E3 recognition) for different retention intervals (RI; E2-E3 lag), using data from all trials (regardless of behavioral responses at E1 and E2). Lines reflect the fits of quadratic trends. Circles mark the peak in the fitted lines. (Also see Supplementary Fig. 1). Qualitatively, the optimal amount of spacing increased as a function of RI. **e,** Subsequent memory (E3 recognition) improved as a function of E1-E2 spacing for images that were successfully recognized at E2 (*β* = 0.10, *p* < 0.001, logistic mixed-effects regression). **f,** Subsequent memory performance (E3 recognition) for images with < 24 h spacing and images with > 24 h spacing (conditional on E2 recognition). Memory was above-chance for both groups (*p*s < 0.001, one-sample *t*-tests) but was significantly greater for images with > 24 h spacing compared to < 24 h spacing (*t* = -2.55, *p* = 0.038, paired *t*-test). **g,** The relationship between spacing and subsequent memory (E3 recognition) was significantly stronger for images successfully recognized at E2 (hit) compared to images not recognized at E2 (miss) (hit: *β* = 0.10, *p* < 0.001; miss: *p* = 0.10, logistic mixed-effects regression; hit compared to miss: *z* = 7.89, *p* < 0.001, *z*-test). Throughout the figure, error bars depict mean ± s.e.m.; dots depict independent participants (*n* = 8); \**p* < 0.05; \*\*\**p* < 0.001.

## RESULTS

### Spaced learning benefits long-term memory

Our primary prediction for behavior was that spacing between the first two exposures (E1-E2 lag) would benefit subsequent memory at the third exposure (E3 recognition). However, prior behavioral work that has considered spacing at long timescales (weeks or months) suggests that memory performance does not monotonically improve as a function of spacing^4^. Rather, as spacing increases, memory benefits first increase, but eventually decrease^6,25^. By some accounts, this non-monotonic function reflects the fact that the benefits of spacing depend on stimuli being recognized when they are re-encountered (recognition at E2 in current study)^10^. From this perspective, the non-monotonic relationship between spacing and memory reflects two competing influences: (i) greater spacing is associated with better memory so long as stimuli are recognized at E2, but (ii) the probability of recognizing a stimulus at E2 decreases with spacing. In the current study, because memory was measured at each encounter (E1, E2, E3), we were able to directly test whether the relationship between E1-E2 spacing (hereafter referred to as spacing) and E3 recognition memory (hereafter referred to as subsequent memory) depended on successful recognition at E2.

An additional nuance to spacing effects is that the optimal amount of spacing (the ‘peak’ in the non-monotonic function) has been shown to increase as a function of the retention interval (RI; here, the RI is the E2-E3 lag)^7,26,27^. For the sake of comparison with this line of work, we report spacing effects as a function of RI for our initial behavioral analyses. However, because our fMRI analyses focus on how E1-E2 spacing influences E1-E2 neural pattern similarity, the RI is not of direct relevance.

Using mixed-effects logistic regression models (see Methods for details), we first tested for relationships between E1-E2 spacing and subsequent memory (hit versus miss at E3), regardless of whether stimuli were successfully recognized at E2 and regardless of behavioral responses at E1. Note that all spacing analyses used the logarithm of the E1-E2 lag intervals (see Methods). Based on prior studies, we predicted a non-monotonic relationship between spacing and subsequent memory, which we tested for as quadratic trends in the logistic regression models. For this first set of analyses, we generated six different models corresponding to RIs ranging from <10 minutes (shortest RI) to > 3 months (longest RI).

Consistent with prior findings^7,28,29^, we found no benefit to spacing for the shortest RI (<10 minutes)—in fact, subsequent memory linearly *decreased* as a function of spacing, with no evidence of a quadratic trend (see Supplementary Table 1 for all statistics). However, for all RIs greater than 10 minutes (the other five models), we observed significant quadratic trends (*p*s < 0.001; Supplementary Table 1). Specifically, as spacing increased, subsequent memory first increased, and then decreased. Qualitatively, the ‘optimal’ amount of spacing (the peak in the quadratic function) increased as a function of the RI (**Fig. 1d** and Supplementary Fig. 1), with the peaks ranging from spacing of ∼1 hour to spacing of several days. Most of the models also had significant negative linear trends (Supplementary Table 1).

We next conducted a separate mixed-effects logistic regression model restricted to stimuli correctly recognized at E2 (E2 = ‘old’ response; hit) and correctly identified as new at E1 (E1 = ‘new’ response; correct rejection); for this model, RI was treated as a nuisance variable (covariate of no interest). We also excluded stimuli for which the RI was < 24 hours because stimuli that were successfully recognized at E2 and then tested less than 24 hours later were effectively at ceiling in terms of E3 memory performance (average hit rate across participants = 0.97). This model again yielded a significant quadratic trend (quadratic term: *β* = -0.01, *p* < 0.001), but in striking contrast to the negative linear trends observed when ignoring E2 recognition (Supplementary Table 1), there was a robust, positive linear relationship between spacing and subsequent memory (*β* = 0.10, *p* < 0.001, logistic mixed-effects regression; **Fig. 1e**). We also directly compared subsequent memory performance (measured as d’) for stimuli that had spacing > 24 hours versus spacing < 24 hours, again restricting analysis to stimuli that were correctly recognized at E2. While performance was well above chance for both conditions (< 24 h: *t* = 9.21, *p* < 0.001; > 24 h: *t* = 5.80, *p* < 0.001, one-sample *t*-tests; **Fig. 1f**), subsequent memory performance was significantly higher for > 24 h spacing compared to < 24 h spacing (*t* = -2.55, *p* = 0.038, paired-samples *t*-test; **Fig. 1f**).

Finally, we also ran a model that was conditionalized on E2 *not* being recognized (E2 = miss; E1 = correct rejection). For this model, we did not observe a significant linear relationship between spacing and subsequent memory (linear trend: *p* = 0.10, logistic mixed-effects regression; **Fig. 1g**). Moreover, the linear relationship between spacing and subsequent memory was significantly stronger when E2 was recognized versus not recognized (*z* = 7.89, *p* < 0.001, *z*-test; **Fig. 1g**). Thus, the benefits of spaced learning were highly dependent on successful recognition at E2.

### Spaced learning strengthens stimulus-specific representations in vmPFC

Having found behavioral evidence that spaced learning benefits subsequent memory, we next assessed whether and how E1-E2 spacing influenced representational similarity across encounters (E1-E2 fMRI pattern similarity). We did this using linear mixed-effects models in which pattern similarity was the dependent measure (see Methods). For these analyses, we did not exclude any trials based on RI. Importantly, however, we did restrict analyses to E1 and E2 trials that were each associated with correct behavioral responses (E1 = correct rejection, E2 = hit) so that any potential relationships between spacing and fMRI pattern similarity were not confounded with behavioral responses. Additionally, to ensure that pattern similarity did not reflect generic cognitive processes, all pattern similarity analyses used a measure of stimulus-specific similarity. That is, for each image we compared ‘within-image’ pattern similarity (E1 and E2 = same stimulus) to ‘across-image’ pattern similarity (E1 and E2’ = different stimuli; **Fig. 2a**). Notably, E2’ images were selected such that they shared behavioral responses with and were presented in the same sessions as E2 (thus approximately matching for spacing; see Methods for details). Stimulus- specific similarity values greater than zero provide positive evidence for a representation of a specific stimulus.

**Fig. 2:**
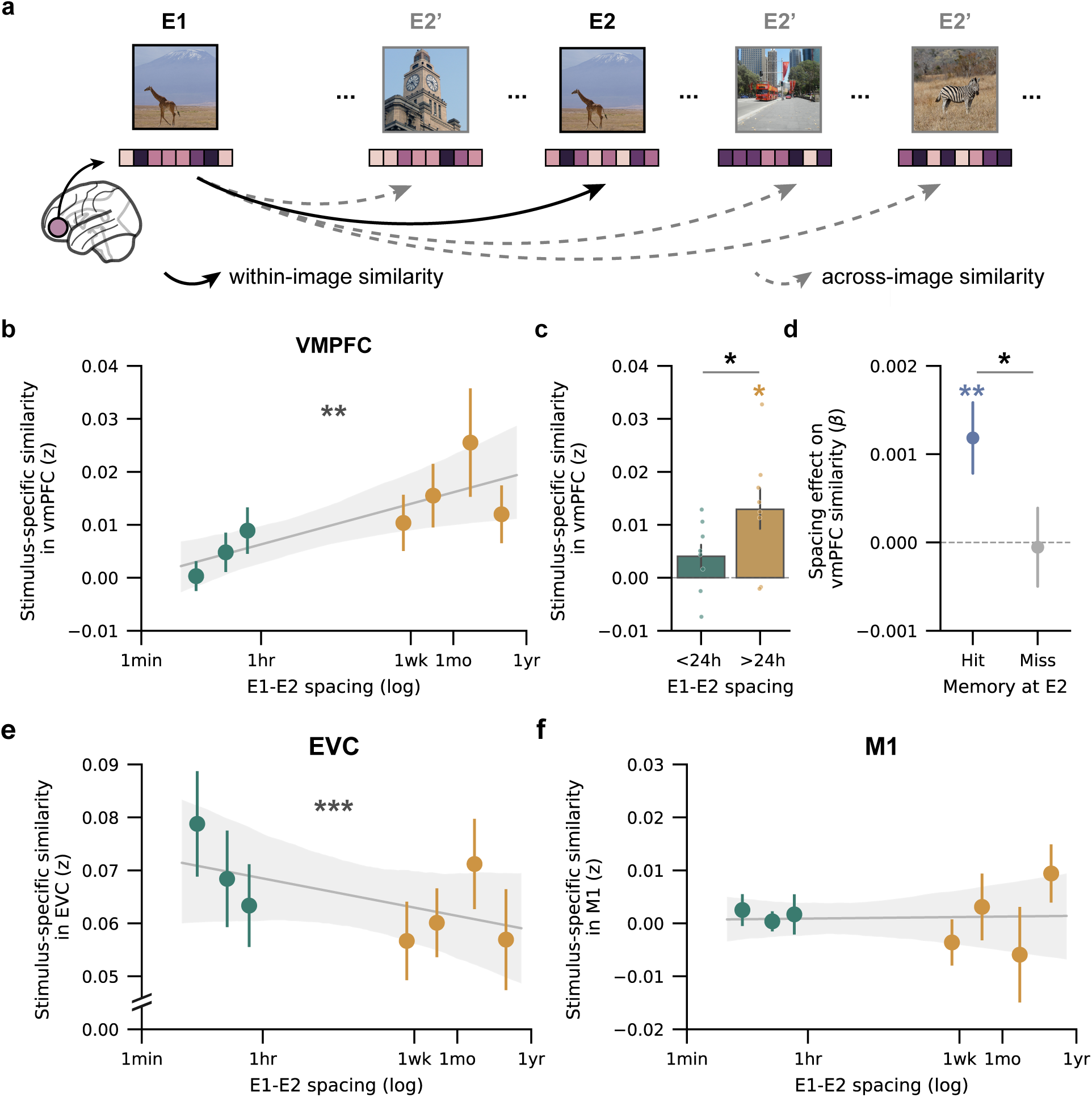
Spaced learning strengthens stimulus-specific representations in vmPFC. **a,** Schematic illustration of stimulus-specific pattern similarity analysis. For each image, we computed pattern similarity scores (z-transformed Pearson correlations) reflecting within-image similarity (E1 and E2 = same stimulus) and across-image similarity (E1 and E2’ = different stimuli). Stimuli used to compute the across-image similarity (E2’ images) were associated with (1) the same behavioral responses as E2 images and (2) the same session as E2 images (E2 session = E2’ session; see Methods for details). For each image, we then computed the difference between within- and across-image similarity, which we refer to as ‘*stimulus-specific’* pattern similarity. Under this difference measure, values greater than zero indicate positive evidence for a representation of a specific stimulus in a given brain region. **b,** Stimulus-specific similarity in vmPFC increased as a function of spacing (*β* = 0.001, *p* = 0.003, linear mixed-effects regression). **c,** Stimulus-specific similarity values in vmPFC were significantly greater for stimuli with > 24 h spacing compared to < 24 h spacing (*t* = -2.82, *p* = 0.026, paired *t*-test). Stimulus-specific similarity in vmPFC did not differ from zero for stimuli with < 24 h spacing (*t* = 1.71, *p* = 0.13, one-sample *t*-test) but was significantly greater than zero for > 24 h spacing (*t* = 3.23, *p* = 0.014, one-sample *t*-test). Dots depict individual participants (*n* = 8). **d,** The relationship between spacing and vmPFC pattern similarity was stronger (more positive) for images successfully recognized (hit) compared to images not recognized (miss) at E2 (hit: *β* = 0.001, *p* = 0.003; miss: *p* = 0.90, linear mixed-effects regression; hit compared to miss: *z* = 2.07, *p* = 0.038, *z*-test). **e,** Stimulus-specific similarity in EVC decreased as a function of spacing (*β* = -0.002, *p* < 0.001, linear mixed-effects regression). **f,** Spacing had no effect on stimulus-specific similarity in M1 (*p* = 0.27, linear mixed-effects regression). Throughout the figure, error bars depict mean ± s.e.m.; \**p* < 0.05; \*\**p* < 0.01, \*\*\**p* < 0.001.

We focused our analyses on three regions of interest (ROIs): (1) vmPFC, given our *a priori* prediction that vmPFC representations would be influenced by spacing^19,21^, (2) early visual cortex (EVC) as a control region that would be sensitive to low-level visual information but would not be expected to contribute to or reflect memory-related effects, and (3) motor cortex (M1) as a control region that would not be expected to be sensitive to visual information or memory-related effects.

Given the positive linear relationship between spacing and subsequent memory that we observed in our behavioral analysis of stimuli that were recognized at E2 (**Fig. 1e**), here we tested for similar linear relationships between spacing and (stimulus-specific) pattern similarity. Intuitively, it might be predicted that greater spacing would be associated with *lower* pattern similarity (more variability). Indeed, EVC exhibited a strong *negative* relationship between spacing and pattern similarity (*β* = -0.002, *p* < 0.001, linear mixed-effects regression; **Fig. 2e**). In other words, EVC similarity decreased as a function of spacing. In sharp contrast, spacing was *positively* related to pattern similarity in vmPFC (*β* = 0.001, *p* = 0.003, linear mixed-effects regression; **Fig. 2b**). That is, the vmPFC representation at E2 exhibited greater similarity to E1 when the E1-E2 lag was longer. Binning stimuli with < 24 h versus > 24 h spacing confirmed that vmPFC pattern similarity did not differ from zero for < 24 h spacing (*t* = 1.71, *p* = 0.13, one-sample *t*-test), but was significantly greater than zero for > 24 h spacing (*t* = 3.23, *p* = 0.014, one-sample *t*-test; **Fig. 2c**). Thus, stimulus-specific representations in vmPFC only emerged when spacing was relatively high (> 24 h). In contrast, EVC exhibited significant pattern similarity for both < 24 h spacing (*t* = 8.55, *p* < 0.001, one-sample *t*-test) and for > 24 h spacing (*t* = 7.99, *p* < 0.001, one-sample *t*-test). Thus, EVC did consistently code for stimulus-specific information, even if these representations were weaker with greater spacing. As expected, we did not observe any evidence that pattern similarity in M1 was influenced by spacing (*p* = 0.27, linear mixed-effects regression; **Fig. 2f**), nor did M1 exhibit stimulus-specific representations for stimuli with < 24 h spacing (*t* = 0.61, *p* = 0.56, one-sample *t*-test) or > 24 h spacing (*t* = -0.18, *p* = 0.86, one-sample *t*-test).

Importantly, we also confirmed that there was no significant linear relationship between spacing and stimulus-specific similarity in vmPFC when images were *not recognized* at E2 (*p* = 0.90, linear mixed-effects regression; **Fig. 2d**). Moreover, the linear relationship between spacing and vmPFC pattern similarity was significantly stronger when stimuli were successfully recognized at E2 compared to when they were not recognized at E2 (*z* = 2.07, *p* = 0.038, *z*-test). Together, these results demonstrate that spaced learning strengthened stimulus-specific representations in vmPFC, but only when stimuli were successfully recognized at E2. These data strongly parallel our behavioral findings: that subsequent memory linearly increased as a function of spacing when stimuli were successfully recognized at E2 (compare **Fig. 2b** to **Fig. 1e**).

It is important to emphasize that conditionalizing the fMRI analyses on successful recognition at E2 avoided potential confounds between spacing and behavioral responses at E2 (namely, as spacing increases there is a lower probability that E2 = hit). Moreover, establishing the relevance of E2 recognition in spacing effects is also of theoretical importance. That said, we also assessed stimulus-specific pattern similarity in vmPFC as a function of spacing for *all trials* (regardless of behavioral response at E2 or E1). This revealed a significant quadratic trend (quadratic term: *β* = -0.001, *p* = 0.04; Supplementary Fig. 2), qualitatively similar to the non-monotonic relationship between spacing and subsequent memory that we observed in the behavioral analyses that included all trials (**Fig. 1d**). Thus, in multiple respects, we observed a strong parallel between pattern similarity in vmPFC and behavioral effects of spaced learning. Importantly, these parallels were particularly evident—or uniquely observable—because we considered a very wide range of timescales (spacing from seconds to months).

### Stimulus-specific similarity in vmPFC predicts behavioral benefits of spacing

We have so far shown that spaced learning (E1-E2 spacing) induced parallel increases in both subsequent memory performance (E3 memory) and stimulus-specific pattern similarity in vmPFC (E1-E2 similarity). We next sought to directly link these behavioral and neural expressions, at the level of individual stimuli. To this end, we used logistic mixed-effects regression models (see Methods), with the dependent variable being subsequent memory and independent variables of pattern similarity and spacing. RI was again included as a covariate and stimuli with RI less than 24 hours were excluded due to ceiling effects in subsequent memory (see *Spaced learning benefits long-term memory*, above). Primary analyses were restricted to stimuli that were successfully recognized at E2, as in the preceding section.

We first tested for a relationship between pattern similarity in vmPFC and subsequent memory and then tested whether this relationship interacted with spacing. The overall relationship between vmPFC pattern similarity and subsequent memory was not significant (*p* = 0.12, logistic mixed-effects model). However, adding an interaction term to the model revealed that the relationship between vmPFC similarity and subsequent memory strongly depended on spacing (*β* = 0.14, *p* < 0.001, logistic mixed-effects regression; **Fig. 3a**). Specifically, the strength of the relationship between vmPFC similarity and subsequent memory increased as a function of spacing. Follow-up tests confirmed a robust positive relationship between vmPFC pattern similarity and subsequent memory when spacing was > 24 h (*β* = 0.81, *p* = 0.002), but not when spacing was < 24 h (*β* = -0.21, *p* = 0.18). This complements findings described above (**Fig. 2b, c**) showing that stimulus-specific representations in vmPFC only emerged as spacing increased. Therefore, the increase in stimulus-specific pattern similarity in vmPFC that emerged with greater spacing was clearly linked to subsequent memory.

**Fig. 3:**
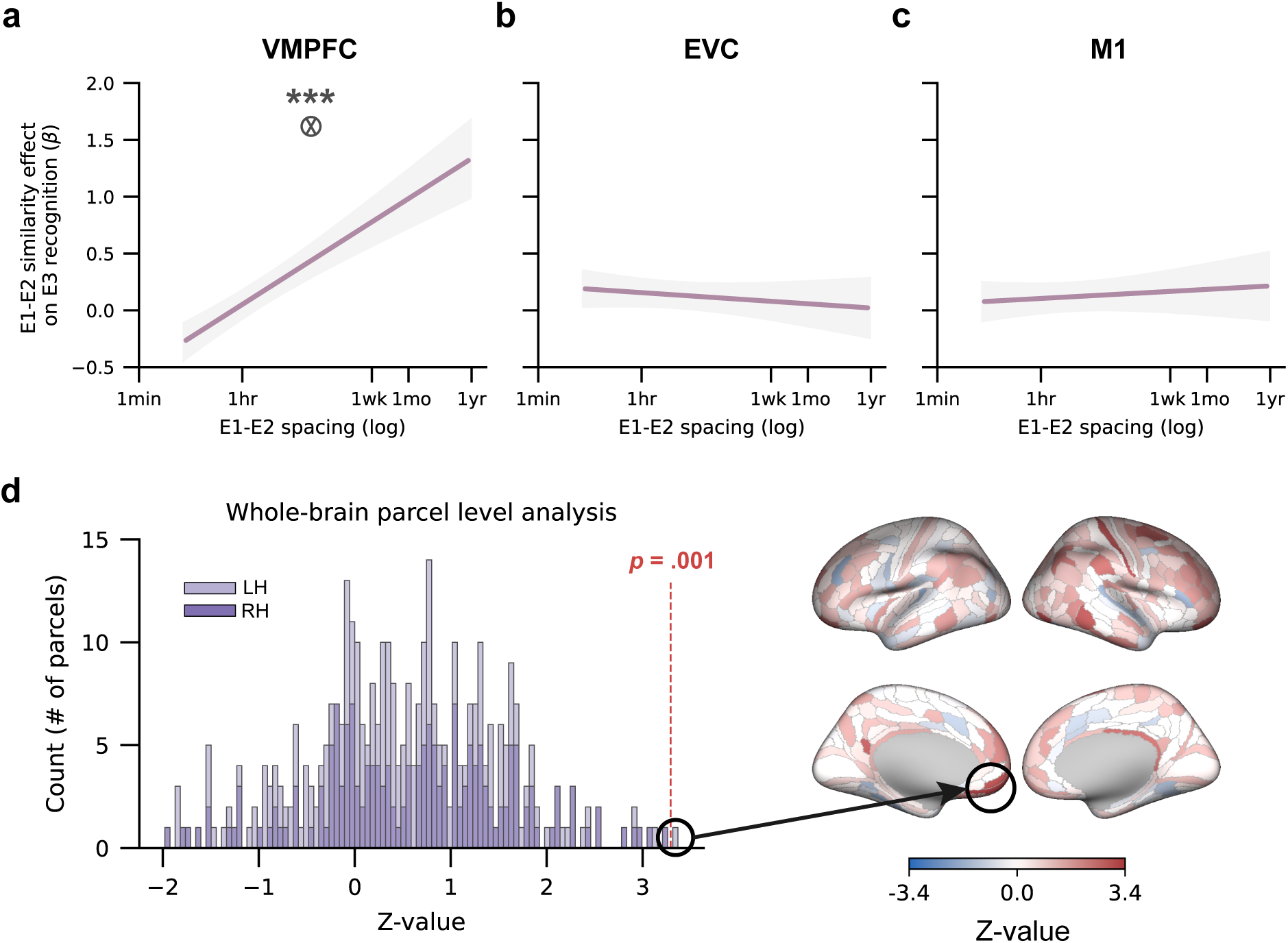
Stimulus-specific similarity in vmPFC predicts behavioral benefits of spacing. **a,** The relationship between vmPFC similarity (E1-E2 similarity) and subsequent memory (E3 recognition) depended on (significantly interacted with) spacing (*β* = 0.14, *p* < 0.001, logistic mixed-effects regression). **b,** The relationship between EVC similarity and subsequent memory did not depend on spacing (*p* = 0.64, logistic mixed-effects regression). **c,** The relationship between M1 similarity and subsequent memory did not depend on spacing (*p* = 0.73, logistic mixed-effects regression). **d,** Exploratory whole-brain analysis to identify cortical regions in which there was a significant interaction between E1-E2 similarity and spacing in predicting subsequent memory. Z values refer to the *z*-scores calculated for the interaction term in a logistic mixed-effects regression model. Using an arbitrary threshold of *p* < 0.001, the only region to show a significant interaction was a region in left vmPFC. LH, left hemisphere; RH, right hemisphere. Throughout the figure, error bars depict mean ± s.e.m.; \*\*\**p* < 0.001; ⊗ indicates spacing × pattern similarity interaction.

We next tested whether pattern similarity in the control ROIs—EVC and M1—was related to subsequent memory. Neither region exhibited an overall relationship between pattern similarity and subsequent memory (*p*s > 0.26), nor did they exhibit an interaction with spacing (*p*s > 0.64). Thus, despite the presence of strong, stimulus-specific pattern similarity in EVC and a significant change in this pattern similarity with spacing (**Fig. 2e**), EVC representations were not predictive of subsequent memory. To further characterize the selectivity of the relationship, we conducted an exploratory whole-brain analysis (including 360 ROIs based on a cortical atlas^30^) to identify regions in which the relationship between pattern similarity and subsequent memory interacted with spacing. Using an arbitrary threshold of *p* = 0.001, this analysis revealed a single region: left vmPFC (**Fig. 3d**). However, as this exploratory analysis only included cortical areas, we also directly interrogated several medial temporal lobe (MTL) regions that are known to be involved in episodic memory^31,32^: hippocampal subfields CA1 and CA2/3/dentate gyrus, entorhinal cortex, perirhinal cortex, and parahippocampal cortex. The relationship between pattern similarity and subsequent memory did not interact with spacing for any of the MTL regions (*p*s > 0.05).

Finally, we tested whether the observed interaction in vmPFC depended on images being successfully recognized at E2, as with the behavioral spacing effects (**Fig. 1g**) and the spacing-dependent increase in vmPFC pattern similarity (**Fig. 2d**). Indeed, the interaction in vmPFC was not significant when the regression analysis was restricted to images that were not recognized at E2 (*p* = 0.55 for interaction term of pattern similarity x spacing on subsequent memory; logistic mixed-effects regression). Thus, at the behavioral and neural levels, the effects of spaced learning depended on successful recognition at E2.

### Stimulus-specific similarity in vmPFC reflects encoding-related processes

Thus far, our findings demonstrate a parallel and direct link between the behavioral benefits of spaced learning and spacing-dependent increases in vmPFC pattern similarity. However, these findings raise the obvious question: *why* does greater spacing increase pattern similarity in vmPFC? The fact that increases in vmPFC similarity depended on E2 recognition might suggest that vmPFC tracked successful retrieval of the original encounter. However, retrieval strength should not increase with greater spacing—it should decrease. Alternatively, many theoretical accounts of spacing effects emphasize the importance of *encoding processes* when a stimulus is repeated^33,34^. Moreover, to the extent that encoding and retrieval are opposing memory states^35–39^, decreases in retrieval strength may, in fact, directly support stronger encoding processes. Thus, an interesting possibility is that vmPFC similarity reflected the (re)encoding of retrieved E1 representations. To address this idea, we tested whether vmPFC similarity was correlated with encoding-related neural processes (and, for comparison, with retrieval-related processes). Leveraging the extensive neuroimaging literature on episodic memory encoding and retrieval, we identified, from independent meta-analyses, an ROI strongly associated with successful memory encoding and an ROI strongly associated with successful recollection (memory retrieval accompanied by contextual information) (see Methods). We then tested, on a stimulus-by-stimulus basis, whether univariate activation in these ROIs at E2 correlated with the degree of stimulus-specific E1-E2 pattern similarity in vmPFC. This was tested using linear mixed-effects regression models with vmPFC similarity as the dependent variable and independent variables including univariate activation (either in the encoding or retrieval ROI) and spacing. These analyses were again restricted to stimuli associated with correct recognition at E2.

Strikingly, higher E1-E2 pattern similarity in vmPFC was predicted by greater E2 activation in the encoding-related ROI (left inferior frontal gyrus; *β* = 0.006, *p* < 0.001, linear mixed-effects regression; **Fig. 4a**) *and* by lower E2 activation in the retrieval-related ROI (left angular gyrus; *β* = -0.003, *p* = 0.040, linear mixed-effects regression; **Fig. 4b**). Importantly, because these models included spacing as a factor, these results indicate that univariate activation in the encoding and retrieval ROIs explained variance in vmPFC pattern similarity above and beyond that explained by spacing alone.

**Fig. 4:**
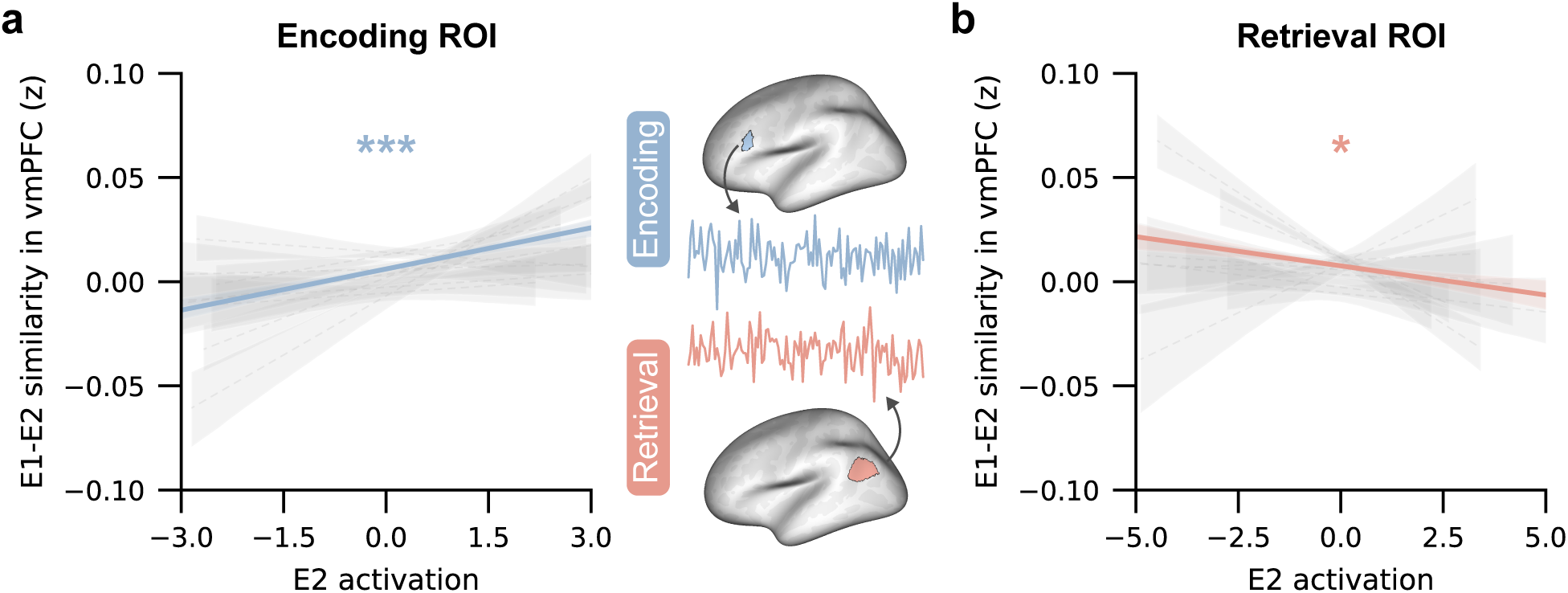
Stimulus-specific similarity in vmPFC reflects encoding-related processes. **a,** There was a positive relationship between stimulus-specific E1-E2 pattern similarity in vmPFC and E2 activation in the encoding ROI (*β* = 0.007, *p* < 0.001, linear mixed-effects regression). **b,** There was a negative relationship between pattern similarity in vmPFC and E2 activation in the retrieval ROI (*β* = -0.003, *p* = 0.04, linear mixed-effects regression). Throughout the figure, gray dashed lines depict independent participants (n = 8); color lines depict group-level relationship; \**p* < 0.05; \*\*\**p* < 0.001.

Exploratory, whole-brain analyses (Supplementary Fig. 3) did not reveal additional ROIs (beyond left inferior frontal gyrus) in which activity positively correlated with vmPFC pattern similarity. However, activity in several additional ROIs (beyond left angular gyrus) was negatively correlated with vmPFC similarity. These regions largely overlapped with the network of brain regions that has been implicated in the recollection of episodic memories^40,41^. Together, these results provide compelling evidence that, for stimuli correctly recognized at E2, greater E1-E2 pattern similarity in vmPFC was related to stronger engagement of encoding-related processes.

## DISCUSSION

Despite over a century of research related to spacing effects in memory, there remains debate about the underlying explanation for why spacing benefits memory. A major limitation to most current theories of spacing effects is that they have not been directly informed by—or constrained by—evidence from neural measures. Relevant neural evidence is limited, at least in part, because it is difficult to continuously measure neural representations over the timescales across which spacing effects operate (hours, days, weeks, or months). Here, we leveraged a unique human fMRI dataset that allowed us to measure neural representations and behavioral effects of spacing at lags that ranged from seconds to many months. By jointly considering these neural and behavioral expressions—and using each measure to understand the other—we were able to gain fundamental new insight into why spacing benefits memory.

An important finding from behavioral studies of spacing effects is that the relationship between spacing and memory is non-monotonic^4^. Based on this finding, leading theories of spacing effects involve two-factor accounts^10,12,16,42^—one factor to explain the ‘rise’ in the spacing function (benefits of spacing) and another factor to explain the ‘fall’ in the spacing function (costs of spacing). Here, we first replicated the classic non-monotonic relationship between spacing and memory when considering data from all trials (without conditionalizing on E2 memory; **Fig. 1d**).

The peak in the observed functions (the optimal amount of spacing) fell somewhere between 1 hour and 1 week, depending on the retention interval (**Fig. 1d** and Supplementary Fig. 1). We then observed a qualitatively similar non-monotonic relationship between spacing and vmPFC pattern similarity (E1-E2 similarity; Supplementary Fig. 2), again with a peak that fell between 1 hour and 1 week. At a broad level, these behavioral and fMRI data establish that the current dataset was extremely well suited to studying spacing effects and highlight the value of considering spacing effects over long timescales.

One of the most prominent explanations for the benefits of spaced learning (the ‘rise’ in the spacing function) is based on the idea of encoding variability^9–11,11,15,25,43,44^. By this account, greater spacing between stimulus exposures leads to more variable encoding of that stimulus (owing to greater change in the encoding context). In turn, a more variably-encoded stimulus has more points of access and, thereby, is more likely to be remembered. Interestingly, we did find that, in early visual cortex (EVC), stimulus-specific representations became markedly less similar (more variable) as a function of spacing (**Fig. 2e**)—a finding that complements other recent evidence^45–47^. However, these EVC effects did not parallel or predict the behavioral benefits of spacing (**Fig. 3b**). In contrast, we found that vmPFC similarity *increased* (became less variable) with greater spacing and, critically, greater vmPFC similarity positively predicted subsequent memory—particularly at long timescales (**Fig. 3a**). Our finding of spacing-related increases in vmPFC similarity is consistent with recent evidence in humans^18^ and rodents^19^ of increases in neural similarity with spacing at short timescales. Our finding that these increases predicted memory aligns with other empirical evidence that neural similarity across stimulus repetitions is positively related to memory^48–50^ and challenges leading theories of spacing effects that emphasize the role of encoding variability. Thus, we were able to identify opposing effects of spacing on representational similarity (increases and decreases in similarity), but we specifically linked the behavioral benefits of spacing to *increases* in neural similarity.

Why would vmPFC similarity (or similarity in any brain region) increase as a function of spacing? One factor that was clearly relevant to the increase in vmPFC similarity was whether events were recognized when they were re-encountered (i.e., at E2). Indeed, we observed a strong linear increase in vmPFC similarity when analyses were conditionalized on successful recognition at E2 (**Fig. 2b-d**), but no evidence of a linear increase in similarity when analyses were conditionalized on failed recognition at E2 (**Fig. 2d**). Importantly, this dissociation parallels what we observed in behavior (**Fig. 1e-g**). Thus, our data strongly support the prominent idea that spacing effects depend on study-phase retrieval^13,14^. That said, while study-phase retrieval was necessary for the benefits of spacing to occur, study-phase retrieval does not, on its own, explain why greater spacing would produce better memory—in fact, the probability of study-phase retrieval should monotonically decrease with greater spacing.

Motivated by theoretical perspectives arguing that encoding and retrieval are opposing neural states^36,51^, we reasoned that decreases in memory retrieval might be beneficial precisely because they allow for greater memory encoding—or, more specifically, *re-encoding* of a retrieved representation. We tested this by correlating, in a stimulus-by-stimulus manner, E1-E2 similarity in vmPFC with the degree of E2 univariate activation in regions of interest that have independently been firmly established as being involved in memory encoding and memory retrieval. Indeed, we found that vmPFC similarity was not only positively correlated with encoding activation, but negatively correlated with retrieval activation. Thus, even though vmPFC similarity critically *depended* on successful retrieval of the original encounter, it was correlated with encoding processes. While this combination may seem contradictory, this is precisely the type of trade-off between opposing influences that is required to explain non-monotonic effects of spacing (in the brain and behavior). Put another way, our findings suggest that memory is likely to benefit when the original experience is successfully retrieved *and* re-encoded. At very short lags, retrieval may be ‘too strong,’ thereby preventing successful re-encoding; at very long lags, even though an encoding state may be ‘strong,’ failure to retrieve the original experience prevents re-encoding. Thus, intermediate lags may be optimal because they allow for a balance between retrieval (of the original exposure) and encoding (of the retrieved memory).

While a re-encoding account provides a parsimonious explanation for our findings—and of spacing effects more generally—a recent computational model suggests a slightly different interpretation that also aligns well with our findings. Namely, Antony et al. argue that variability triggers the abstraction of similarities across stimulus exposures^26^. That is, when a stimulus is re-encountered after a relatively long delay, the encoding context is more likely to be different and this difference triggers error-driven learning that strengthens common elements across encounters at the expense of unique elements (i.e., abstraction of similarities). The key point is that this account explains spacing effects in terms of increased neural similarity when events are spaced across time. However, an abstraction account does make a unique prediction about memory for contextual information—including memory for when in time each encounter occurred (temporal memory). Specifically, an abstraction account predicts that, with relatively long spacing, retrieval of the original encounter should actively weaken temporal memory for individual encounters. On the one hand, this prediction is challenged by existing behavioral^13^ and fMRI evidence^50^ indicating that retrieval of past encounters actually enhances temporal memory. On the other hand, it is notable that temporal memory does not necessarily benefit from spaced learning^50^. Thus, in future work it will be of interest to further characterize the relationship between spaced learning and contextual memory (including temporal memory) as understanding this relationship will help refine theoretical accounts.

The fact that our findings specifically implicate vmPFC in representing event similarity over time is notable in light of the extensive literature demonstrating vmPFC involvement in the integration of information across encoding events^52,53^ and in the formation of schemas^54,55^. In particular, vmPFC is thought to support the encoding of new events into existing schematic representations^53,54^—a function which resembles the re-encoding account proposed above. Other studies have also implicated vmPFC in the retrieval of remote memories^21–24,56,57^ and in memory consolidation^24,58,59^. Notably, some of this evidence specifically relates to vmPFC activation increasing as a function of the age of a retrieved memories^58,59^—with the idea being that memories ‘move’ to (or emerge within) vmPFC over time. In contrast, we show that when a stimulus is re-encountered after a relatively long lag, this can actually increase the similarity *to the original experience*. Thus, our findings are not readily consistent with an account where vmPFC representations only emerge over long timescales. That said, other theories of consolidation allow for the possibility that vmPFC representations are formed immediately, but the reliance on these representations changes over time^24^.

In summary, by considering learning events that were spaced from seconds to many months apart, we show that the behavioral benefits of spaced learning are strongly paralleled—and predicted by—the similarity with which vmPFC represented stimuli across exposures. Moreover, through a number of complementary analyses, we show that the benefits of spaced learning—in brain and behavior—reflect a balance between retrieval and encoding processes. Namely, whereas retrieval of an original encounter is a necessary condition for the benefits of spaced learning, it is also important that retrieved information is re-encoded. From this perspective, non-monotonic relationships between spacing and memory can be explained by trade-offs between these two computations.

## METHODS

### Overview

This study reports findings using the Natural Scenes Dataset (NSD; http://naturalscenesdataset.org). The NSD is a large-scale fMRI dataset in which participants performed a continuous recognition task on thousands of color natural scenes over the course of 30–40 high-resolution fMRI (7T) scan sessions. The results in this study are based on data from all of these sessions, from all participants who took part in the NSD study. For a detailed description of the dataset, including methods involved in preprocessing the fMRI data, please see the original data publication^20^. Below we outline the specific methods relevant to the current study.

### Natural Scenes Dataset

#### Participants

Eight participants from the University of Minnesota community participated in the NSD study (two self-identified males and six self-identified females; age range 19–32 years). All participants were right-handed with no known cognitive deficits nor color blindness and with normal or corrected-to-normal vision. Participants were naïve to the design of the NSD experiment and were not involved in the design or planning of the study. Informed written consent was obtained from all participants before the start of the study, and the experimental protocol was approved by the University of Minnesota Institutional Review Board.

#### Design and procedure

A detailed description of the experimental design has been reported in the original data publication^20^. Briefly, participants performed a continuous recognition task in which they reported whether the current image had been seen at any previous point in the experiment (‘old’) or if they had not encountered it before (‘new’). For each participant, the experiment was split across 40 scan sessions in which 10,000 distinct color natural scenes would be presented three times spaced pseudo-randomly over the course of the entire experiment. Each scanning session consisted of 12 runs (750 trials per session). Each trial lasted 4 s and consisted of the presentation of an image for 3 s and a following 1-s gap (**Fig. 1a**). Participants were able to respond during the entire 4-s period and were also permitted to make multiple responses per trial in cases where they changed their mind. As trials with multiple responses potentially captured more complex cognitive operations related to decision making, we opted to exclude those trials from all analyses.

Participants completed up to 40 scan sessions each and were scanned approximately once a week over the course of 10 months (**Fig. 1b**; the first scan session for each participant corresponds to Day 0). Four of the participants completed the full set of 40 NSD scan sessions. Due to constraints on participant and scanner availability, two participants completed 30 sessions and two participants completed 32 sessions. Accordingly, each participant viewed a total of 9,209- 10,000 unique images across 22,500-30,000 trials.

#### Stimuli

Images used in NSD were taken from the Microsoft Common Objects in Context (COCO) database^60^. A total of 73,000 unique images were prepared with the intention that each participant would be exposed to 10,000 distinct images (9,000 unique images and 1,000 shared images across participants) three times each across the 40 scan sessions. Image exposures were pseudo-randomly distributed across sessions over the course of almost a year. The presentation structure was determined in advance and fixed across participants so that difficulty of the recognition task was roughly similar across participants. Distribution of image presentations was controlled to ensure that both short-term and long-term re-exposures were probed. To provide a sense of the overall experimental design, the mean number of distinct images shown once, twice, and all three times within a typical session is 437, 106, and 34, respectively.

As the current study focused on how the spacing between the first two exposures to an image influenced recognition at the third exposure, we only considered images that were presented all three times across the entire experiment. Here, we labeled each exposure based on its presentation order (i.e., the first exposure is E1, the second exposure is E2, and the third exposure is E3). Of critical interest, the spacing between the first two exposures (E1-E2 lag) ranged from 4 seconds to 302 days (see Fig. 1c for distribution of the E1-E2 spacing for each participant).

#### fMRI data acquisition and preprocessing

MRI data was collected at the Center for Magnetic Resonance Research at the University of Minnesota. Imaging was performed on a 7T Siemens Magnetom passively-shielded scanner with a single-channel-transmit, 32-channel-receive RF head coil. Functional images were acquired using whole-brain gradient-echo echo-planar imaging (EPI) at 1.8-mm resolution and 1.6-s repetition time.

Details of the preprocessing of anatomical and functional data are provided in the original data publication^20^. Briefly, functional data were preprocessed by performing one temporal resampling to correct for slice time differences and one spatial resampling to correct for head motion within and across scan sessions, EPI distortion, and gradient non-linearities. Informed by the original publication, the current study used the 1.0-mm volume preparation of the functional time-series data and “version 2” of the NSD single-trial betas.

### Data Analysis

Based on the observation that stimuli that were successfully recognized at E2 and then tested less than 24 hours later (E2-E3 lag < 24 hours) were effectively at ceiling in terms of E3 memory performance (average hit rate across participants: 0.97), all analyses in the current study related to E3 memory only excluded stimuli for which the E2-E3 lag was > 24 hours.

#### Behavioral data analyses

We first separately tested for relationships between spacing (E1-E2 lag) and subsequent memory (E3 memory) for different retention intervals (RI; E2-E3 lag), regardless of whether stimuli were successfully recognized at E2 (and regardless of behavioral responses at E1). To do so, we generated six different models corresponding to RIs ranging from <10 minutes (shortest retention interval) to > 3 months (longest retention interval). For each RI, we performed the mixed-effects logistic regression analyses that predicted subsequent memory from spacing using both linear and quadratic fits. Specifically, we used a mixed-effects logistic regression model that predicted subsequent memory (hit = ‘old’ response, miss = ‘new’ response) from spacing while including (controlling for) several additional factors. These added factors of no interest included the lag between the beginning of the first trial in the experiment and the first exposure (i.e., E1 onset), the retention interval (i.e., E2-E3 lag), and false alarm rates of sessions in which each exposure occurred. All models were constructed with random intercepts for each participant. Because memory is observed to abide by an exponential rule rather than linear time^61^, all temporal lag information (i.e., E1 onset, E1-E2 lag, and E2-E3 lag) was quantified by expressing time intervals in seconds and transforming these intervals with the natural logarithm.

To directly test whether the relationship between spacing and subsequent memory (and fMRI pattern similarity) depended on successful recognition at E2, we first ran analyses restricted to stimuli correctly recognized at E2 (E2 = hit) and correctly rejected at E1 (E1 = ‘new’ response; correct rejection) and then ran analyses conditionalized on E2 not being successful recognition at E2 (E2 = miss; E1 = correct rejection).

#### Regions of interest (ROIs) definition

To probe whether the spacing showed a modulation of stimulus-specific representations, a region-of-interest (ROI) analysis was performed. Motivated by prior evidence implicating vmPFC in episodic memory across long timescales^21–24^ and by recent evidence of spacing effects on neural similarity in the rodent medial prefrontal cortex^19^, this analysis was focused on the bilateral ventromedial prefrontal cortex (vmPFC), along with two control ROIs: bilateral early visual cortex (EVC) and bilateral motor cortex (M1). All cortical ROIs were drawn from the surface-based Human Connectome Project multimodal parcellation (HCP-MMP) atlas of human cortical areas^30^. An additional exploratory analysis was conducted throughout the whole brain using all parcels available in the HCP-MMP atlas. Based on the established role of the medial temporal lobe (MTL) in human memory, we also repeated certain analyses in a set of subregions there. These MTL ROIs included bilateral CA1, CA2/3/dentate gyrus, entorhinal cortex, perirhinal cortex, and parahippocampal cortex, all manually drawn on the high-resolution T2 images obtained for each participant.

To probe specific hypotheses concerning encoding/retrieval-related processes, a priori ROIs were chosen based on past work. Specifically, an ROI representative of encoding-related effects was identified using the global peak coordinates supporting encoding success from an independent meta-analysis on the subsequent memory effect^62^. We then projected the coordinates to the HCP-MMP atlas and used the delimiting parcel as the encoding ROI. Similarly, for an ROI representative of retrieval-related effects, we used the peak coordinates supporting recollection success from another independent meta-analysis of episodic memory retrieval^63^ and identified the delimiting parcel in the HCP-MMP atlas as the retrieval ROI.

#### Pattern similarity analyses

Pattern similarity was calculated as the Pearson correlation between activity patterns evoked during different image exposures for each ROI. Correlations were z-transformed (Fisher’s z) before further analyses were performed. To avoid potential contamination from BOLD signal autocorrelation, all pattern similarity analyses were performed by correlating activity patterns for stimuli across run (i.e., correlations were never performed within the same scanning run).

For our primary analyses related to pattern similarity between E1 and E2, of critical interest was the stimulus-specific pattern similarity. Specifically, for each image (‘target’), we compared ‘within-image’ pattern similarity (E1 and E2 = same stimulus) to ‘across-image’ pattern similarity (E1 and E2’ = different stimuli; Fig. 2a). The E2’ images used to compute across-image similarity were chosen from the same set of sessions (but different scanning runs) as the target image’s E1 and E2, thus controlling for differences in spacing. Further, E2’ images were chosen such that they had the same memory outcomes at the first two exposures as the target image. For target images with multiple possible E2’ images based on the criteria above, the median value of the across-image similarity scores was used. The across-image similarity was then subtracted from within-image similarity to yield a stimulus-specific similarity measure of E1-E2 similarity for each image.

Unless indicated otherwise, our main fMRI analyses were restricted to stimuli that were associated with correct behavioral responses at both E1 and E2 (i.e., E1 = ‘new’ responses, E2 = ‘old’ responses) so that any potential relationships between spacing and neural pattern similarity were not confounded with behavioral responses.

### Statistical analyses

Behavioral and fMRI data were analyzed using a combination of paired *t* tests and mixed-effects regression models. Relationships between spacing, stimulus-specific pattern similarity and subsequent memory were tested with mixed-effects regression models. For all mixed-effects regression models, we used the participant as a random effect and other variables as fixed effects. All *t*-tests were two-tailed. A threshold of *p* < 0.05 was used to establish statistical significance for all analyses unless otherwise specified. fMRI analyses were corrected for multiple comparisons when applicable.

## Acknowledgement

We thank D. L. Hintzman and members of the Kuhl, DuBrow, and Hutchinson labs for fruitful discussions. This work was supported by NIH-NINDS 2R01NS089729 to B.A.K.

## Data availability

Data from the NSD is publicly available at http://naturalscenesdataset.org.

## Code availability

Code for analyses in this study will be made available upon publication at https://github.com/futingzou.

## Competing interests

The authors declare no competing financial interests.

## Supplementary Information

**Supplementary Fig. 1.**
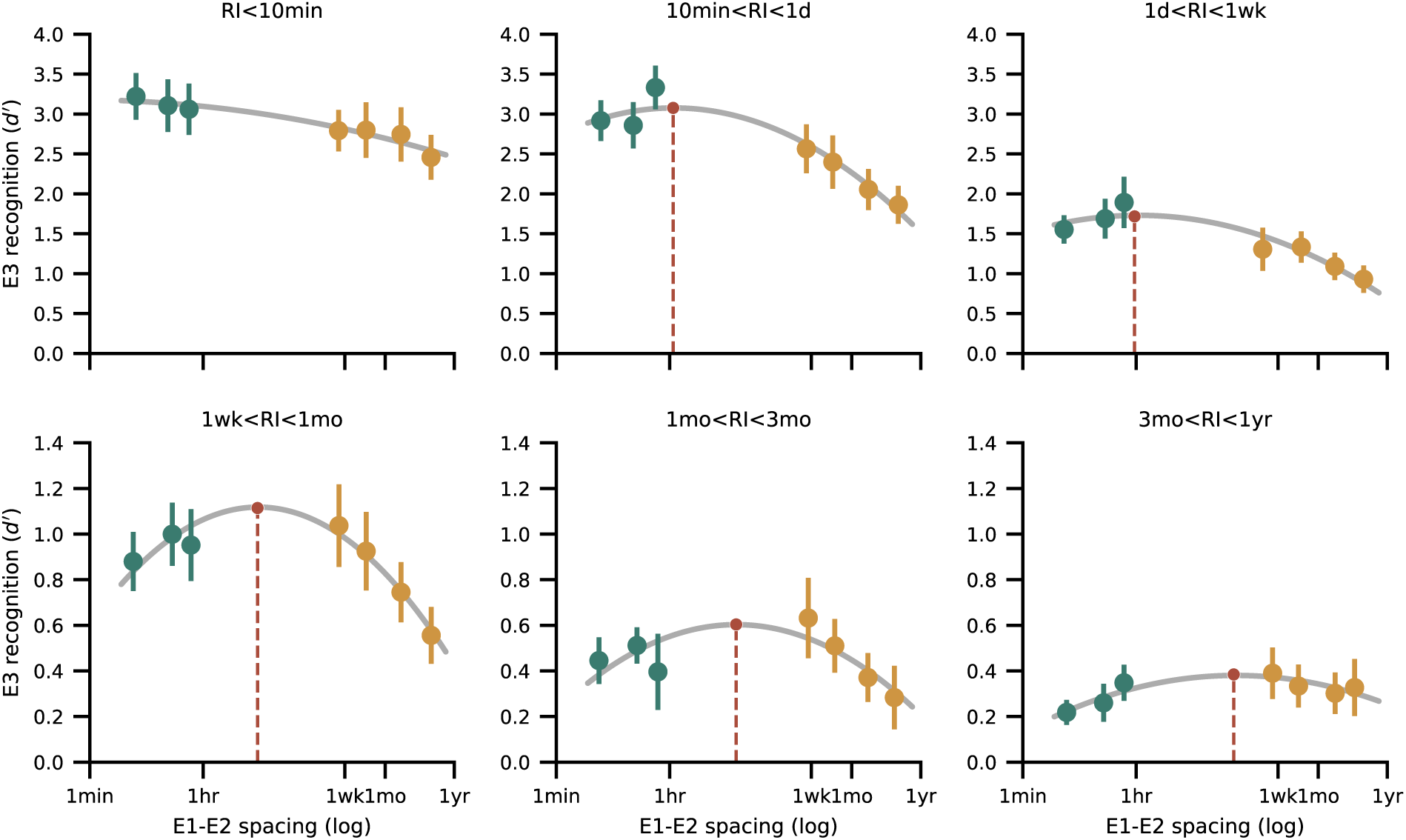
Relationships between spacing and subsequent memory (recognition accuracy) for different retention intervals (RIs) for all trials (regardless of whether stimuli were successfully recognized at E2, and regardless of responses at E1).

**Supplementary Fig. 2.**
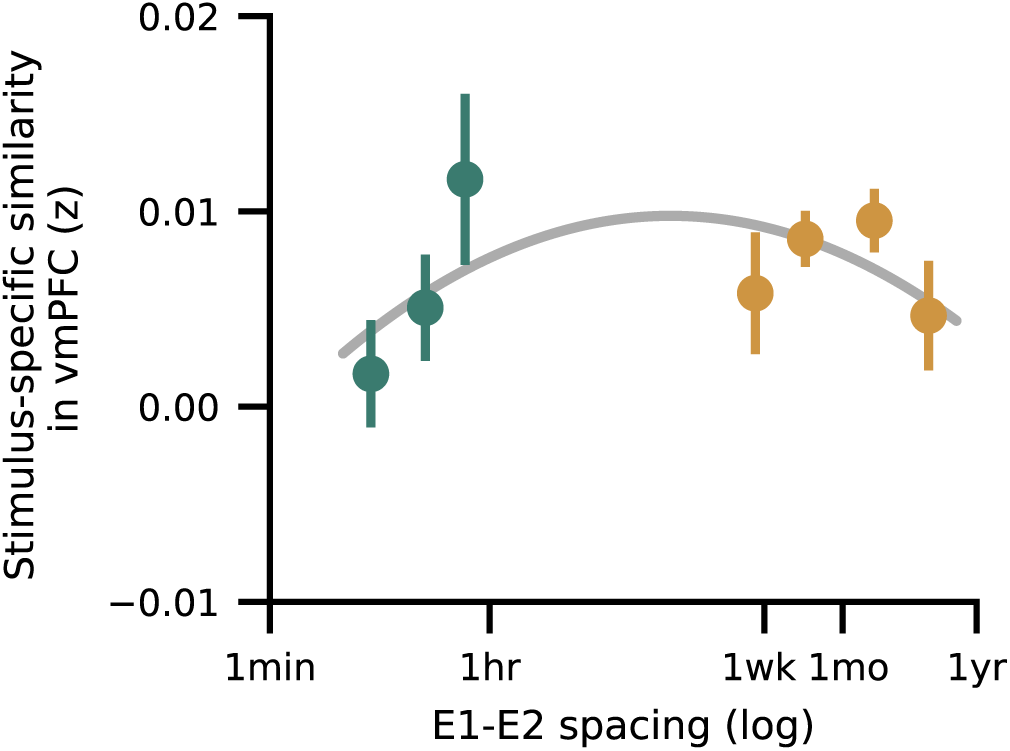
Relationship between spacing and stimulus-specific pattern similarity in vmPFC for all trials (regardless of whether stimuli were successfully recognized at E2, and regardless of responses at E1).

**Supplementary Fig. 3.**
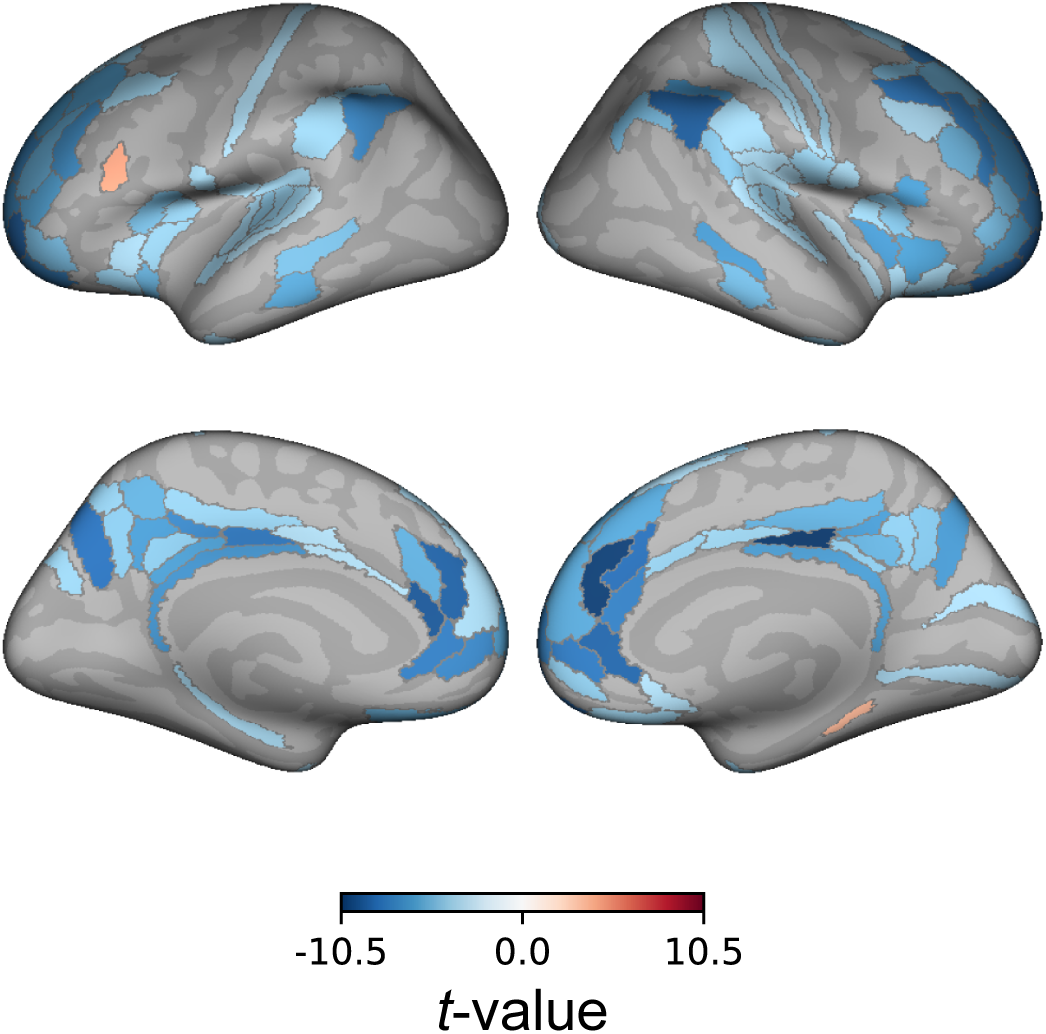
Exploratory whole-brain analysis. Linear mixed-effects regression analyses were performed across the whole brain to test for relationships between E2 univariate activation in cortical regions and E1-E2 pattern similarity in vmPFC. Warm colors reflect regions in which E2 univariate activation positively correlated with E1-E2 similarity in vmPFC. Cool colors reflect regions in which E2 univariate activation negatively correlated with E1-E2 similarity in vmPFC.

**Supplementary Table 1.** Statistics for relationships between E1-E2 spacing and subsequent E3 memory for different retention intervals (E2-E3 lag) using both linear and quadratic trend analyses for all trials (i.e., regardless of behavioral responses at E1 and E2). To compare the fits of models for each RI, we used ANOVA to test whether adding the quadratic term of spacing led to significantly better prediction of subsequent memory relative to models without the quadratic term.

| Retention interval (RI) | Linear fits | Quadratic fits (quadratic term) | Model comparison |
| --- | --- | --- | --- |
| RI<10min | $\beta = -0.130$<br>$p < 0.001^{***}$ | $\beta = -0.004$<br>$p = 0.425$ | $\chi^2 = 0.623$<br>$p = 0.430$ |
| 10min<RI<1d | $\beta = -0.157$<br>$p < 0.001^{***}$ | $\beta = -0.017$<br>$p < 0.001^{***}$ | $\chi^2 = 25.146$<br>$p < 0.001^{***}$ |
| 1d<RI<1wk | $\beta = -0.113$<br>$p < 0.001^{***}$ | $\beta = -0.023$<br>$p < 0.001^{***}$ | $\chi^2 = 53.789$<br>$p < 0.001^{***}$ |
| 1wk<RI<1mo | $\beta = -0.045$<br>$p < 0.001^{***}$ | $\beta = -0.022$<br>$p < 0.001^{***}$ | $\chi^2 = 70.151$<br>$p < 0.001^{***}$ |
| 1mo<RI<3mo | $\beta = -0.012$<br>$p = 0.079$ | $\beta = -0.013$<br>$p < 0.001^{***}$ | $\chi^2 = 25.021$<br>$p < 0.001^{***}$ |
| 3mo<RI<1yr | $\beta = 0.016$<br>$p = 0.007^{**}$ | $\beta = -0.009$<br>$p < 0.001^{***}$ | $\chi^2 = 13.965$<br>$p < 0.001^{***}$ |

## Notes

### Competing Interest Statement

The authors have declared no competing interest.

